# Progressive oxygenation of developing leaves directs morphogenesis

**DOI:** 10.1101/2025.09.21.677629

**Authors:** Gabriele Panicucci, Vinay Shukla, Viktoriia Voloboeva, Leonardo Jo, Kees van Kollenburg, Sara Buti, Laura Dalle Carbonare, Francesco Licausi, Daan A. Weits

## Abstract

Oxygen availability underpins energy production in multicellular organisms, yet internal oxygen gradients arise in both plants and animals. Plants sense these variations via the PLANT CYSTEINE OXIDASE branch of the N-degron pathway, which regulates the stability of key transcription factors. Originally linked to metabolic control, this pathway recently emerged as a development regulator. While the shoot apical meristem was shown to be hypoxic, the oxygen dynamics of organs originating from this low-oxygen niche remain unknown. Here we show that developing leaves form a spatiotemporal oxygen gradient that is sensed through the oxygen sensing machinery. This pathway integrates local oxygen availability to regulate leaf morphogenesis: early hypoxia restricts cell expansion, while subsequent distal-to-proximal oxygenation enables specialized cell fates acquisition. Our findings reveal that oxygen acts as a positional cue in normal growth, guiding developmental trajectories. Our work highlights opportunities to harness oxygen gradients or sensing to direct plant form and function.

## Introduction

Molecular oxygen leads a double life in plants. On the one hand, oxygen’s primary recognized contribution to plant physiology is serving as the final electron acceptor in mitochondrial aerobic energy production. On the other hand, oxygen also functions as a signaling molecule that influences the stability of key transcription factors that trigger both stress responses and developmental processes. This regulatory role is mediated by the activity of PLANT-CYSTEINE-OXIDASES (PCOs), which use oxygen as a co-substrate to oxidize proteins harboring an exposed N-terminal Cysteine ^1–3^. This post-translational modification acts as a tag for degradation through the N-degron proteolytic pathway, *de facto* rendering these proteins oxygen-labile ^4^. Scarcity of molecular oxygen, which leads to the stabilization of PCO substrates, can occur acutely as a result of exogenous calamitous events such as flooding, or chronically, as observed in meristematic tissues such as the shoot apical meristem (SAM) ^5,6^. Perturbation of this endogenous hypoxic state, for instance through hyperoxia treatments, reduces leaf initiation rates, suggesting that hypoxia is not merely tolerated in the SAM but instead potentially plays an active role in maintaining meristematic function ^7^. Group VII ETHYLENE RESPONSE FACTORS (ERFVII), LITTLE ZIPPER 2 (ZPR2) and VERNALIZATION 2 (VRN2), oxygen-labile PCO substrates, are stable in the hypoxic meristematic niche and contribute to the regulation of root architecture, meristem activity, vernalization and cell expansion ^7–10^.

While the hypoxic nature of the SAM and its functional implications have been shown previously, the oxygen dynamics of organs originating from this hypoxic niche are so far unexplored. Leaves start their developmental track as a bulge of proliferative cells on the periphery of the hypoxic SAM. As the primordium emerges, proliferation is gradually confined to the basal portion of the leaf and eventually stops; in its place, differentiation proceeds from the apical portion of the leaf towards the base, leading to the formation of specialized cell types ^11^. While auxin and cytokinin hormones are known to regulate cell proliferation and expansion during leaf development, the exact mechanisms orchestrating the distal-to-proximal progression of this process remain incompletely understood ^12,13^.

To fulfill their roles as the primary photosynthetic organs and regulators of gas exchange, water use and thermoregulation, leaves build several specialized structures and cell types across their development. Crucial to their role as a source organ, chlorophyll is produced and accumulated in differentiated mesophyll cells. Gas exchange, necessary for photosynthesis, is provided by the development of guard cells through the commitment of protodermal cells to the stomatal cell lineage ^14^. Mechanical resistance across the leaf surface is guaranteed by the interdigitation of epidermal cells, which during cellular expansion switch from isotropic to anisotropic growth creating convex and concave lobes ^15^. Trichomes, originating from de-differentiated epidermal cells confer an additional layer of protection against predation and increase water retention on the leaf surface ^16,17^.

Leaf morphology varies drastically across different species and developmental stages, and leaf shape is also closely tuned with its environment ^18^. For example, larger leaves harvest more light, but are more prone to tearing by wind, while serrated leaves deter herbivory and improve gas exchange. The coordinated balance between proliferation and differentiation shapes the final morphology of the leaf. When these phases of leaf development are disturbed, for instance through ectopic expression of the meristem regulator *STM* in emerging leaves, differentiation is delayed in favor of a prolonged proliferative phase ^19^. This leads to an increased number and greater depth of serrations and lobes along the leaf margin.

Mature leaves exchange gasses with the surrounding atmosphere, resulting in an internal oxygen content that is close to atmospheric level; leaves are in fact designed to optimize gas exchange through specialized cell types and air spaces ^20^. The final morphology and cellular structure of mature leaves depend, however, on processes that are initiated in the early stages of leaf development, in close proximity to the hypoxic SAM ^21^. In this work, we set out to define the dynamics of internal oxygen content during primordia emergence and to elucidate the potential role of oxygenation in the overall development of leaves.

## Results

### Early leaf development occurs under hypoxic conditions, followed by progressive tip-to-base oxygenation

Leaf primordia emerge from the SAM, a region previously shown to be chronically hypoxic (3.6-8.4 kPa), and this hypoxic state is necessary for proper organ initiation ^7^. Fully mature leaves instead exhibit an oxygen level close to that of the surrounding air. However, little is known as to when leaves oxygenate as they grow out from the hypoxic niche that encloses the shoot apical meristem. To investigate oxygenation dynamics in early leaf development, we first imaged activation of a known hypoxia responsive gene (HRG) *PCO1*, and found that not only the meristem, but also emerging leaf primordia activate low-O_2_ signaling (**Fig.1a**). This suggests permanence of the meristematic hypoxic microenvironment in the early stages of leaf development. We therefore set out to measure actual oxygen content in emerging primordia through O_2_ microprofiling (**Fig.1b**). Our measurements revealed that leaf oxygenation happens gradually during development, rather than abruptly, suggesting that low O_2_ signaling attenuates as leaves emerge from the shoot apex. The gradual nature of oxygenation led us to question whether the entire leaf is oxygenated uniformly, or rather spatially distinct regions of the leaf might experience different oxygenation dynamics. While microsensor measurements offer a reliable quantification of organ- or tissue-level oxygen tension, they are not suitable for examining oxygen levels across an entire organ. In fact, each measurement damages cells and likely creates oxygen entry points. To circumvent these issues, we resorted to the HRG-based reporter line *pPCO1:GUS-GFP* to visualize differences in low oxygen signaling across the full developmental track of a newly formed leaf. GFP Fluorescence showed a clear tip-to-base deactivation of hypoxia-signaling, with *pPCO1:GFP* activity being confined to the basal region in maturing leaves (**Fig.1c**). GFP signal quantification across leaf length, used as a proxy for developmental stage, clearly showed a decrease in hypoxia-signaling as leaves mature (**Fig.1d**). GUS staining of leaves 7 and 19 also showed a clear pattern of distal-to-proximal oxygenation, suggesting that this gradient is consistent along the vegetative phase (**Fig.1e**). Hypoxia in developing tissues has previously been shown to regulate developmental processes, raising the question of whether it may also act as a developmental cue during leaf formation.

**Figure 1.**
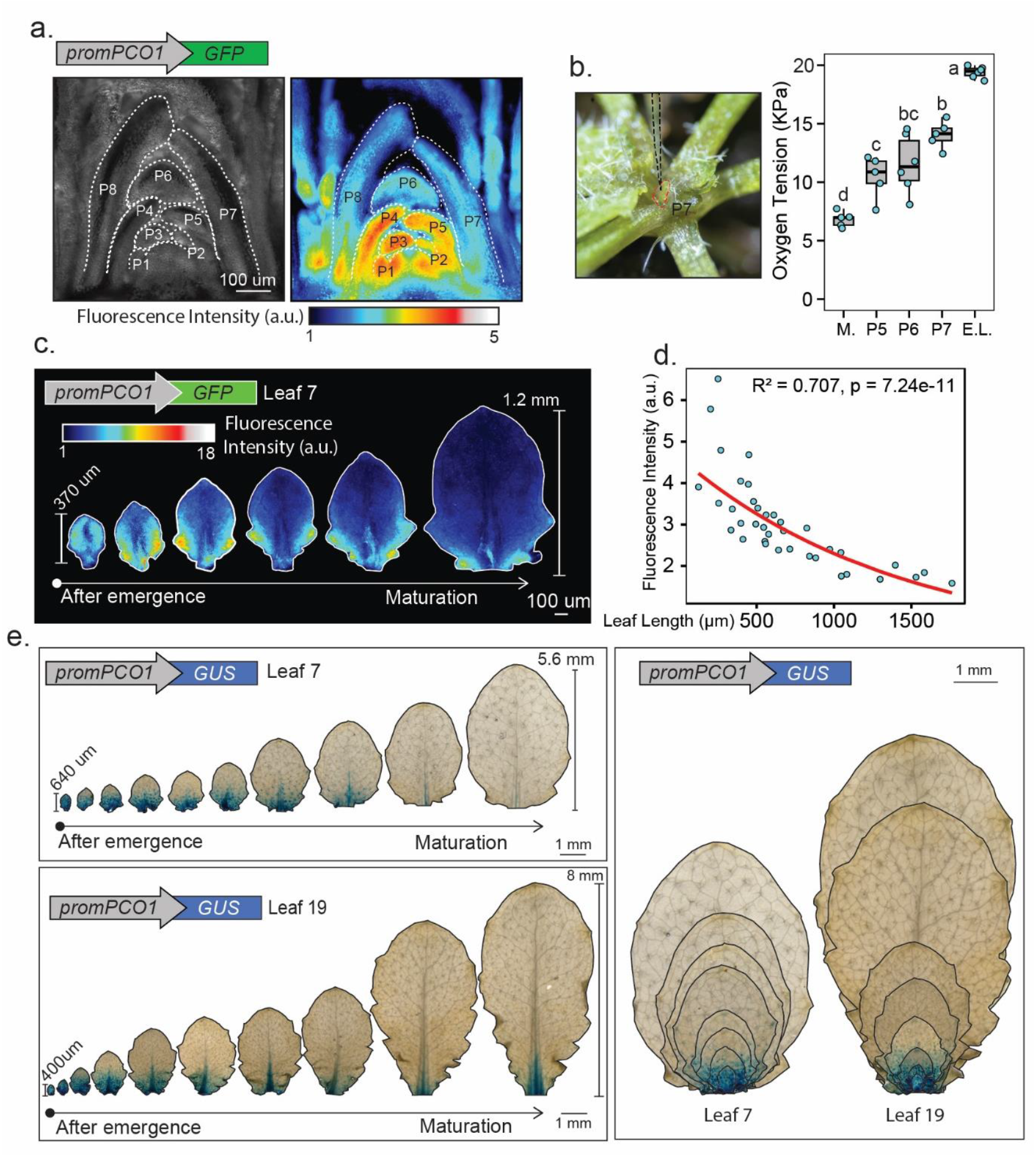
Leaves originate as an hypoxic organ and progressively oxygenate tip-to-base. **(a)** Longitudinal vibratome sections of *Arabidopsis thaliana* shoot apical meristem and emerging leaf primordia. GFP signal visualized with royal look-up-table. Primordia (P) indicated by dashed lines and numbered starting from the most recently emerged (P1). **(b)** Oxygen microprofiling of the meristem dome (M.), emerging leaf primordia (P5,6,7) and fully emerged leaves (E.L.). Microsensor position indicated via a dashed black line, outline of a P7 leaf primordia in red. **(c)** Imaging of GFP signal across *promPCO1:GFP* developing leaves 7 ± 1, from emergence until maturation. Signal visualized with royal look-up-table. **(d)** Quantification of GFP signal intensity in *promPCO1:GFP* developing leaves in relation to the size of their major axis. Red line shows the exponential fit obtained by linear regression of log-transformed GFP signal against length, back-transformed to the original scale. Model fit statistics are indicated on the plot (R^2^ and p-value for the slope). **(e)** GUS staining images of *promPCO1:GUS* leaf 7 and 19 across development.

### Oxygen distribution affects leaf morphology

In order to investigate the possibility that oxygen distribution acts as a developmental signal in leaf development, we investigated whether modified oxygen atmospheres would result in altered leaf morphology. To this end, plants with 10 visible leaves were transferred into either oxygen-depleted (4% O_2_, v/v) or oxygen-enriched (40% O_2_, v/v) artificial atmospheres (**Fig. 2a**). They were maintained under these conditions for 8 days and subsequently returned to ambient air (21% O_2_ v/v). Leaves that developed during the treatment completed their growth and were then scored for morphological traits. Leaves formed under hypoxic conditions were not affected in size, but displayed increased complexity, characterized by deeper serrations in the proximal region (**Fig. 2b**). By contrast, leaves produced under hyperoxia showed a marked reduction in both major and minor axis length, which resulted in an overall reduced size. Serration depth was also diminished, although not significantly, and this likely reflected the overall reduction in leaf size (**Fig. 2b**). In order to uncouple developmental responses from potential pleiotropic effects of hyperoxia, we next opted for an attenuation of hypoxia-responses through the use of an estradiol-inducible *PCO4* construct (**Fig. 2c**). *PCO4* induction during leaf development altered leaf area, primarily as a result of a reduced minor axis length (**Fig. 2d**). Serration depth was also reduced concomitant with a reduced leaf size. Taken together, these data suggest that the persistence of hypoxic conditions in lieu of progressive leaf oxygenation affects final shape to a more serrated, less solid leaf. Conversely, alleviation of low oxygen or low-oxygen signaling during the initial phases of leaf development led to stunted growth, indicating that an hypoxic environment might be needed for the early stages of leaf development in order for leaves to reach their optimal size.

**Figure 2.**
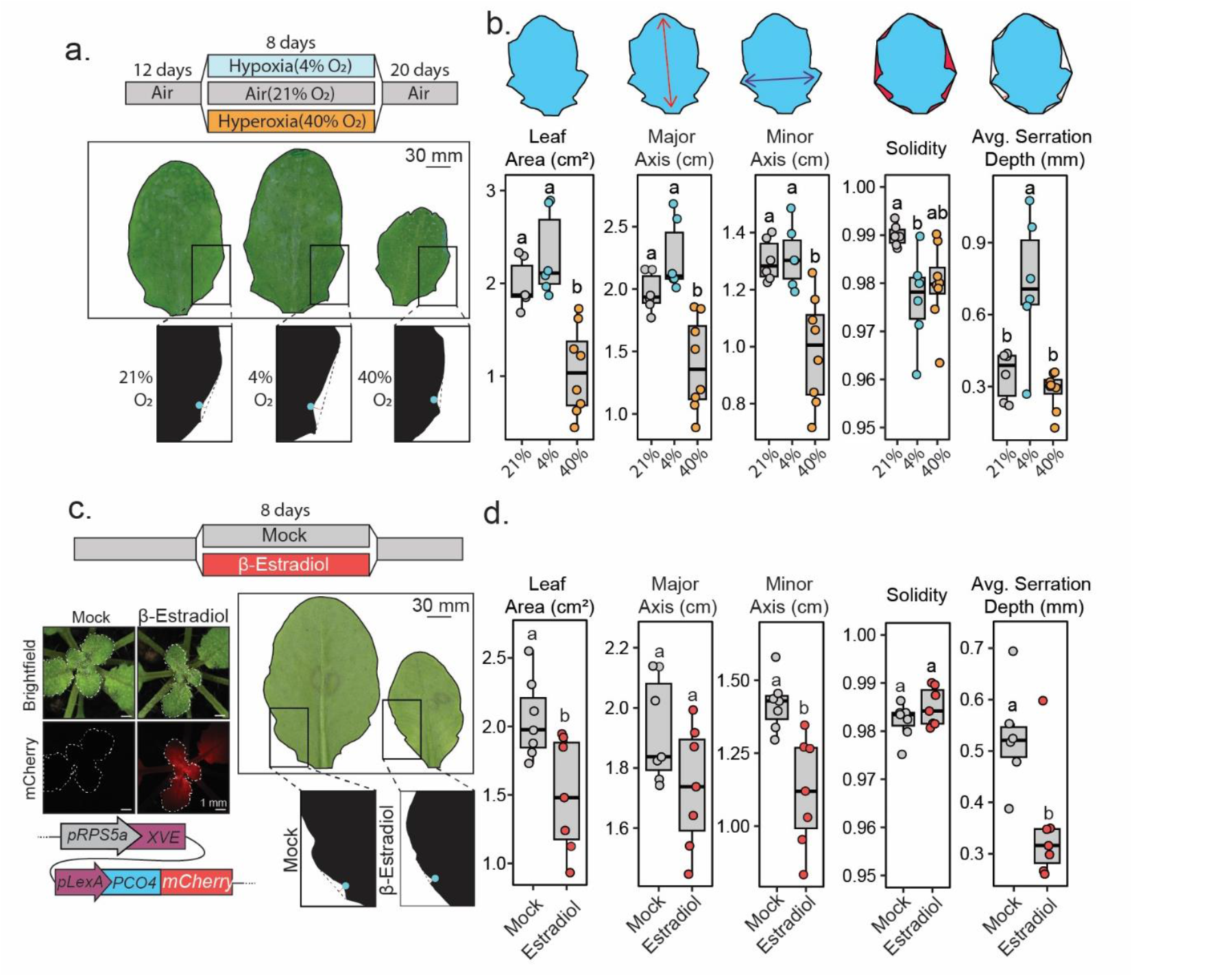
Manipulation of oxygen and oxygen signaling affects leaf shape and size. **(a)** Final shape of fully developed leaf 7 ±1 subjected to 8 days of either 4% or 40% oxygen v/v atmospheres. Serration depth indicated on leaf silhouettes. **(b)** Morphological quantification of leaf shape and size. Solidity is calculated as the ratio between leaf area and the area of its convex hull. Average serration depth is calculated based on the depth of the first two proximal serrations visible on the leaf blade. A schematic representation of each leaf trait parameter is shown above. One-way Anova followed by Tukey post hoc test for multiple comparison was performed to assess statistical significance, with letters indicating statistically different groups (p value < 0.05). **(c)** Final shape of fully developed leaf 7 ± 1 subjected to 8 days of *pRPS5a:XVE-pLEXA:PCO4-mCherry* induction. Scheme of the construct and induction visualization, via mCherry fluorescence, are also reported. Serration depth indicated on leaf silhouettes. **(d)** Morphological quantification of leaf shape and size. Student t-test was performed to assess statistical significance. Letters indicate a statistically significant difference (*p* value < 0.05)

### Role of the PCO branch of the N-degron pathway in leaf development

Plants sense acute and chronic hypoxia via the Cys-branch of the N-degron pathway. To investigate if this oxygen sensing pathway plays a role in integrating oxygen distribution into leaf morphology, we next set out to examine the effects on leaf shape of different oxygen signaling mutants that exhibit either higher or lower N-degron pathway substrates levels (**Fig.3a**). We found that the *4pco* mutant, which constitutively signals hypoxia, produces distinctly more complex leaves from the wildtype (**Fig. 3b**). Leaves of *4pco* mutants are also smaller and rounder, but most strikingly, leaf-margin analysis revealed that average serration depth was markedly enhanced, resulting in a drastic decrease in solidity. Since the *4pco* mutant signals hypoxia even in normoxic tissue and serration depth was also increased in hypoxic conditions, this suggests that the inability to sense leaf oxygenation leads to a strongly altered, more complex, leaf shape. Considering that the phenotype observed in *4pco* did not match the previously published phenotypes of either *ZPR2* or *VRN2* overexpression ^8,22^, we hypothesized that the observed *4pco* leaf complexity traits might depend on persistent stabilization of ERFVII transcription factors during leaf development. In order to test this hypothesis, we genetically removed *RAP2*.*2, RAP2*.*12* and *RAP2*.*3* in the *4pco* background (*4pco3rap*) through CRISPR/Cas9 (Supplementary Data, **Fig. S1**). We also studied leaf phenotypes in a pentuple *erfvii* mutant, where all ERFVIIs were genetically inactivated. The *4pco3rap* mutant displayed a slightly smaller than wild-type leaf surface area and a complete rescue of *4pco* margin-specific phenotypes, suggesting that leaf-shape related features observed in *4pco* are indeed dependent on ectopic stabilization of ERFVII (**Fig.3a,b**). Next, we set out to investigate how the differences observed in final leaf morphology are linked to differences in cellular morphology across the different mutants. To assess this, we examined the size of abaxial epidermal cells in *4pco, erfvii* and *4pco3rap* fully developed leaves. We examined both distal and proximal region, since these exhibit different oxygen dynamics during growth (**Fig.3c**). The *erfvii* mutant displayed markedly increased epidermal cell size and decreased cell number in the distal region and a minor decrease, although not significant, in cell number in the proximal region. Only a minor increase in cell size was observed in *4pco3rap*, indicating that the stabilization of other PCO substrates was largely sufficient to rescue the increased cellular size observed in *erfvii* leaves. Interestingly, the *4pco* mutant did not display differences in epidermal cell size and number in either the proximal or distal region, indicating that the observed leaf shape phenotype is not due to altered patterns of cell expansion. Taken together, our data shows that low oxygen signaling in early stages via *erfvii* regulates cell expansion, while permanence of hypoxic signaling in later stages leads to increased leaf complexity.

**Figure 3.**
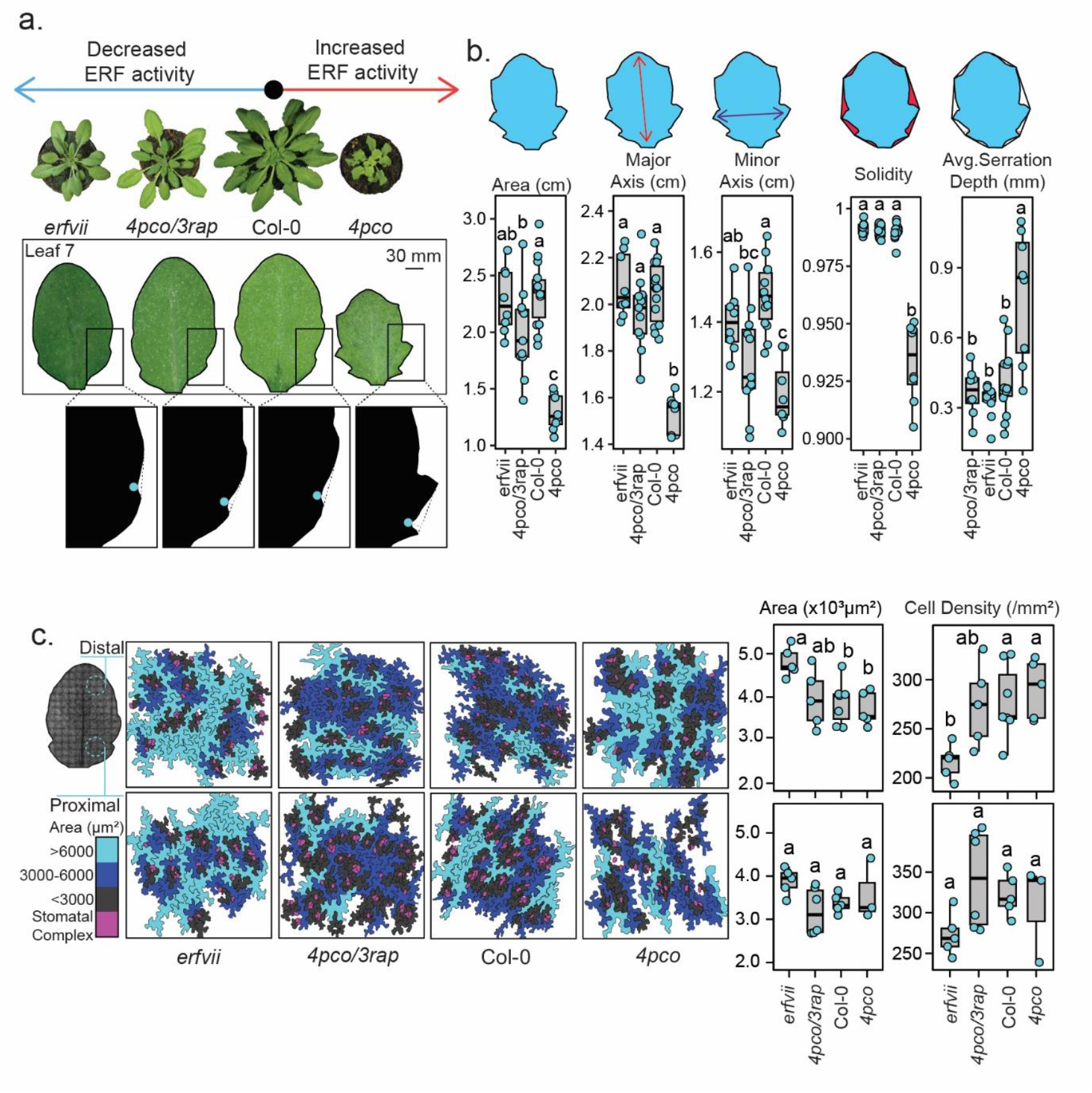
The PCO branch of the N-degron pathway affects leaf development. **(a)** Final shape of fully developed leaf 7 in different N-degron mutants. Genotypes are ordered based on the predicted activity of ERFVII transcription factors in each respective mutant. Serration depth indicated on leaf silhouettes. **(b)** Morphological quantification of leaf shape and size. Solidity is calculated as the ratio between leaf area and the area of its convex hull. Average serration depth is calculated based on the depth of the first two proximal serrations visible on the leaf blade. A schematic representation of each leaf trait parameter is shown above. One-way Anova followed by Tukey post hoc test for multiple comparison was performed to assess statistical significance, with letters indicating statistically different groups (p value < 0.05). **(c)** Cellular phenotype of different N-degron mutants in the distal and proximal regions of fully developed leaf 7. Segmented epidermal cells are colored based on area (cyan for cells larger than 6000 µm2, blue for cells between 3000 and 6000 µm, grey for cells smaller than 3000 µm). Stomatal complexes and stomal lineage cells are indicate in magenta. Epidermal cells size and density are shown as box plots, where each point indicates the average epidermal cell size or density across one leaf. One-way Anova followed by Tukey post hoc test for multiple comparison was performed to assess statistical significance, with letters indicating statistically different groups (p value < 0.05).

### Transcriptome changes underlying the interplay between oxygen sensing and leaf development

In order to understand how the dynamic oxygenation of leaves is connected at the molecular signaling level to oxygen sensing, we conducted an RNA-seq analysis. Here, we compared the leaf transcriptome of the *4pco* mutant with wildtype leaves at different stages of development. These were chosen based on the oxygenation status, with stage 1 and 3 representing fully hypoxic and fully oxygenated leaves, respectively, and stage 2 representing a partially oxygenated leaf (**Fig. 4a**). Differential gene expression was analyzed using a generalized linear model with explanatory factors for genotype, developmental stage, and their interaction. We confirmed the hypoxic signature of the examined leaves by analyzing the expression pattern of the 49 known hypoxia core genes^23^; of these, 38 were upregulated regardless of developmental stage. Among those, 15 showed a significant interaction effect, suggesting a stage-specific layer of regulation on their expression (**Fig. 4b**). Furthermore, among genes which exhibited a significant interaction or genotype effect, we selected those that were associated with leaf development GO terms (**Fig. 4c**). This lead us to identify several misregulated genes involved in cell-fate acquisition such as regulators of trichome patterning and stomata development. For instance, *MYC4*, which was strongest induced in *4pco* stage 3, acts upstream of SPEECHLESS and FAMA to negatively regulate stomatal index, whereas *SOL1* is repressed across all stages of *4pco* and is required for fate transitions within the stomatal lineage ^24,25^. *TRICHOMELESS 2 (TCL2)* and *TOPOISOMERASE 6B* (*TOP6B)* show opposite genotype effects; the former is induced in *4pco* and is a strong repressor of trichome formation, while the repressed gene *TOP6B* is required for endoreduplication and the *top6b* dwarfed mutant shows less trichomes and less lobey pavement cells ^26,27^. *STICHEL* (*STI*) shows both a genotype and interaction effect and regulates trichome branching ^28^. Additionally, we identified numerous differentially expressed genes involved in chlorophyll biosynthesis, repair and overall chloroplast organization. showing genotype, interaction or both effects. A small subunit of RuBisCo, *DEG24*, whose mutation correlates with stunted growth and pale leaves, also shows severe downregulation. Strikingly, few well described meristem maintenance genes were also affected including *SHOOTMERISTEMLESS* (*STM*) and *BARELY ANY MERISTEM 2* (*BAM2*) ^29,30^. Together, the *4pco* transcriptome data shows misregulation of genes involved in the specification of specialized leaf cells, chlorophyll biosynthesis and an expansion of meristem maintenance gene expression, suggesting that constitutive hypoxia signaling delays leaf differentiation.

**Figure 4.**
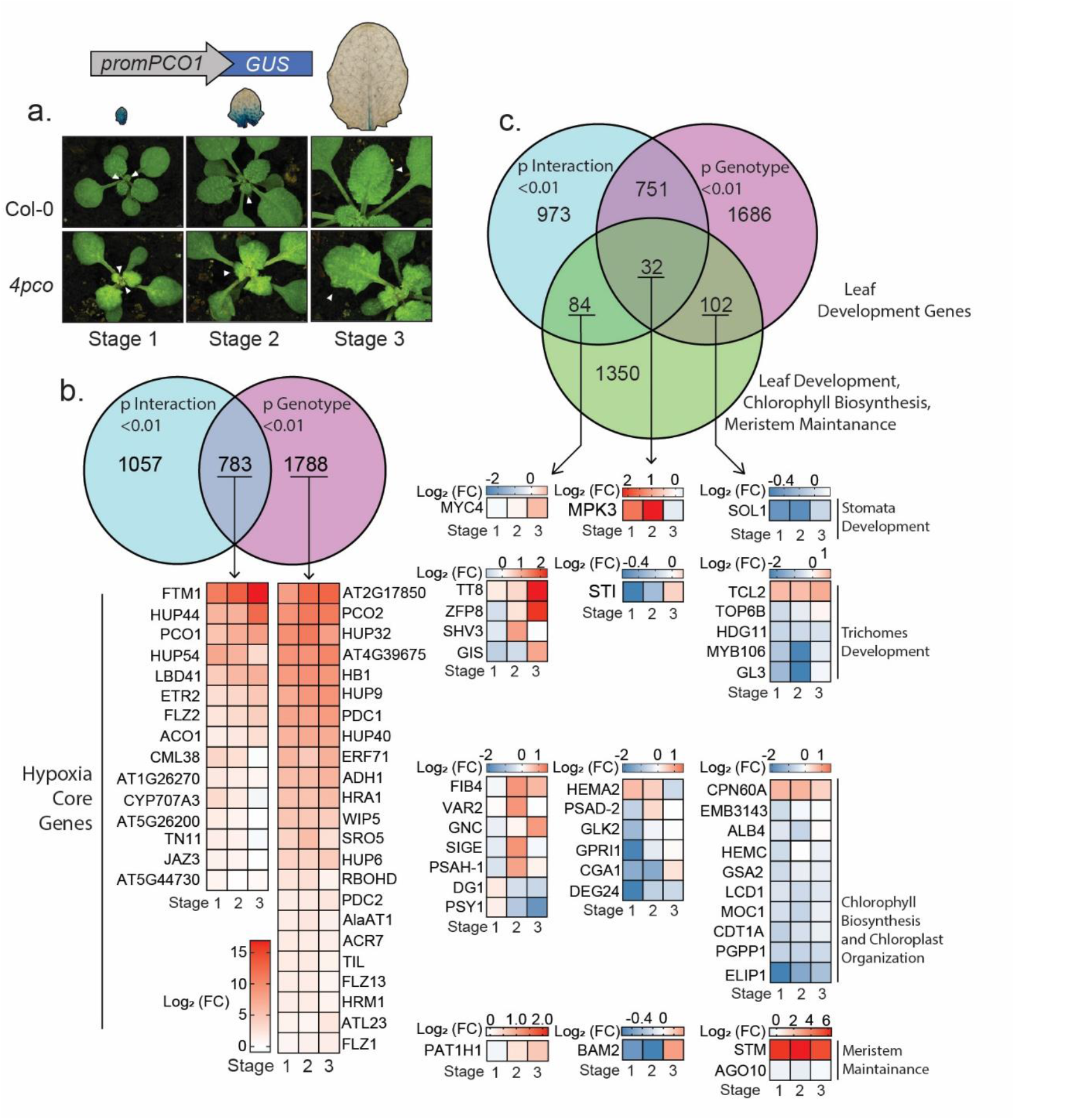
Transcriptome analysis on *4pco* reveals interplay between oxygen sensing and leaf development. **(a)** Example of sampled Col-0 and *4pco* leaves 7±1 according to their hypoxic signature, inferred by activation of *promPCO1:GUS*. **(b)** List of core hypoxia genes that exhibit either a significant interaction effect or both an interaction and genotype effect. Heatmaps show Fold Change across stages 1 to 3 in a log_2_ scale. **(c)** Venn diagrams show the overlap between genotype, interaction effect and leaf developmental genes. List of selected leaf development and meristem maintenance genes that exhibit either a significant interaction effect, genotype effect or both interaction and genotype effect. Heatmaps show Fold Change across stages 1 to 3 in a log_2_ scale.

### Cellular phenotyping reveals how oxygen sensing shapes leaf morphogenesis

In order to assess how the apparent imbalance in cell fate acquisition identified by our transcriptome analysis affected the formation of specialized cell types, we quantified leaf adaxial trichomes and employed a combination of Differential Interference Contrast (DIC) microscopy and cellular segmentation to examine epidermal cells and stomal complexes in oxygen sensing mutants. As reported, the shape of epidermal pavement cells starts isotropic and becomes progressively more complex during leaf development, exhibiting their characteristic lobey puzzle shape in mature leaves ^15^. Interestingly, we observed that the wildtype displays a different pavement cell lobeyness in distal and proximal parts of the leaf (**Fig5a)**. Pavement cells in *4pco* are similar to wildtype cells in the basal part of the leaf, while cells in the distal region never reach the degree of complexity seen in the wildtype (**Fig.5a**). This suggests that the inability of *4pco* to perceive distal leaf oxygenation might result in simpler pavement cells. Furthermore, the number of stomatal complexes in the *4pco* distal and proximal part of the leaf was severely reduced (**Fig.5b**) and a small number of these complexes exhibited irregular, single cell stomatal precursor rather than fully formed guard cells (**Fig.5c**). Considering that the number of irregular stomata only amounted up to 15% of all stomatal complexes, the differences observed in the total number of stomatal complexes in *4pco* are largely due to failure to initiate stomal fate. Both the decrease in lobeyness and in stomatal density are rescued in the *4pco3rap* mutant, suggesting that the impairment in cell fate acquisition is dependent on ERFVII stabilization. Additionally, trichome density was found to be reduced in the *4pco* mutant (**Fig.5d**) and partially rescued in the *4pco3rap* mutant, providing evidence of yet another specialized cell type that is affected by constitutively-on low oxygen signaling. Finally, altered chlorophyll associated transcripts prompted us to investigate its biosynthesis in developing leaves. Wildtype leaves display an inverse-correlation between hypoxia-signaling and chlorophyll fluorescence, suggesting antagonistic regulation (**Fig. 1 and Fig 5e**, Supplementary Data **Fig. S2a,b**). This is supported by *4pco* chlorophyll fluorescence quantification, which was found to be lower than that of wildtype leaves across development (**Fig.5e**). Collectively, cellular phenotyping supports a model in which disrupting oxygen sensing impairs both the acquisition of specialized cell types and chlorophyll biosynthesis.

**Figure 5.**
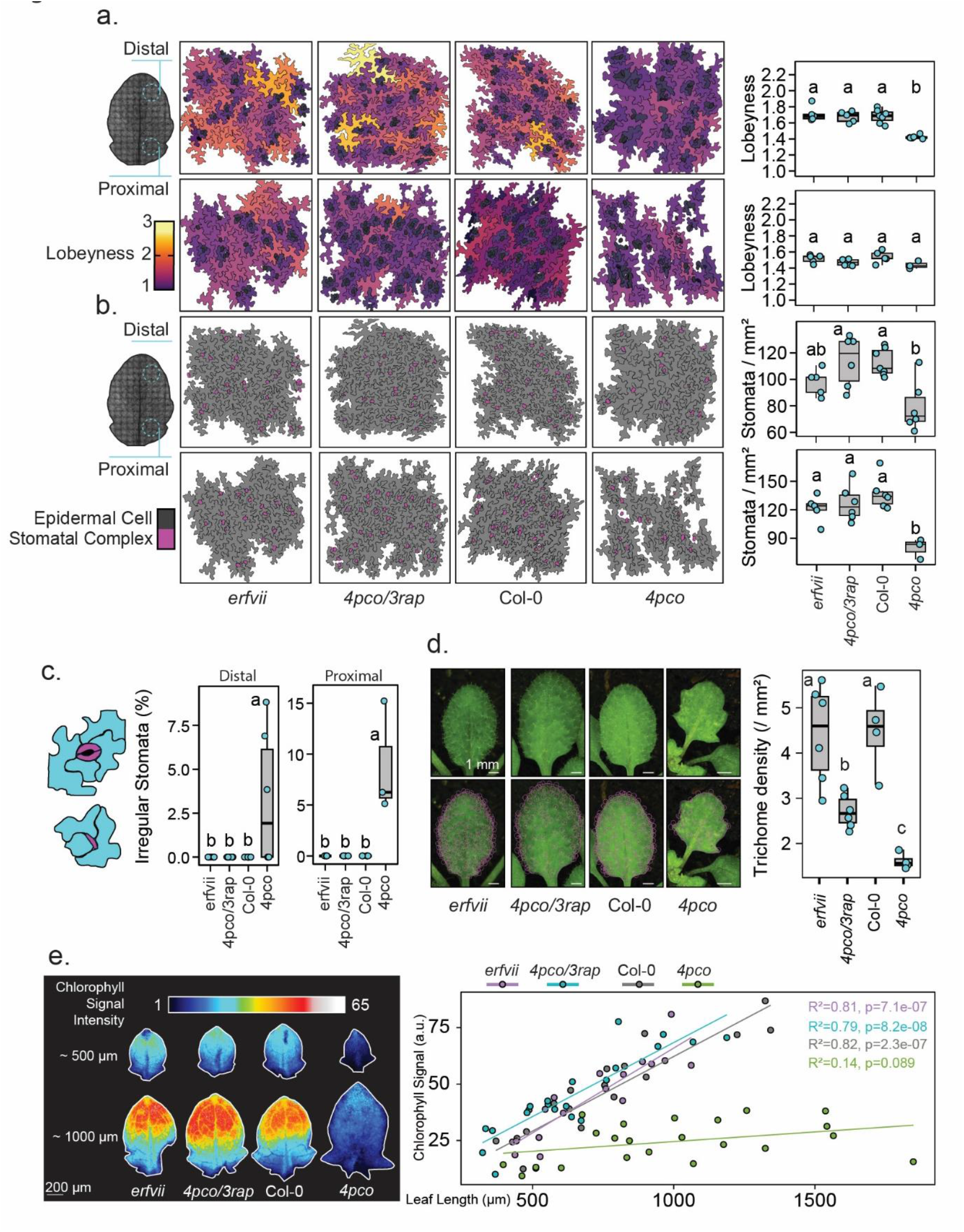
N-degron mutants are impaired in producing specialized leaf cell types. **(a)** Leaf 7 proximal and distal epidermal cells of N-degron mutants colored based on their lobeyness, defined as the ratio between their perimeter and the perimeter of their convex hull. On the right, quantification of average lobeyness across genotypes. Each point represents the average lobeyness of epidermal cells across one leaf. **(b)** Stomatal complexes found in distal and proximal region of N-degron mutants leaf 7. Stomatal density was calculated as number of recognizable pairs of guard cells per mm^2^ of leaf surface. **(c)** Percentage of irregular stomatal complexes identified in proximal and distal regions of various N-degron mutants. (**d**) Number of trichomes identified in various N-degron mutants per mm^2^ of leaf surface. For **(a), (b), (c)** and **(d)** one way Anova followed by Tukey post-hoc test was used to assess statistical differences, with letters indicating statistically different groups (p value < 0.05). **(e)** Chlorophyll autofluorescence measured in leaf 7 ± 1, visualized through royal look-up-table. On the right, quantification of chlorophyll signal intensity over leaf length, with fitted linear regression for each genotype. Model fit statistics are indicated on the plot (R^2^ and p-value for the slope).

### Molecular mechanisms underpinning leaf development are targets of oxygen sensing

Next, we investigated how oxygen sensing mechanisms link to developmental genes previously identified in our RNA-seq analysis. Since the knock-out of *RAP2*.*2,RAP2*.*3* and *RAP2*.*12* in *4pco3rap* was sufficient to largely rescue the *4pco* phenotype, we reasoned that genes required for cell-fate acquisition during leaf development might be regulated by ERFVII. To test this hypothesis, we measured the expression of identified *4pco* misregulated genes in isolated mesophyll protoplasts transformed with constitutive stable RAP2.3. First, the induction of several hypoxia core genes attested that the introduction of RAP2.3 was sufficient to activate hypoxic responses in isolated protoplasts (**Fig.6a**). Then, we found that in accordance with our transcriptome data, negative regulator of trichome formation *TCL2* was activated by RAP2.3, while positive regulators of endoreduplication and trichome formation and branching *STI* and *TOP6B* were repressed (**Fig.6a**). Surprisingly, bHLH transcription factors *MYC3* and *MYC4*, negative regulators of stomatal development, were repressed by RAP2.3, while they were mildly induced in our *4pco* transcriptome data. Trichome regulators *TCL1, GLABRA3* (*GL3*), and *ZINC*-*FINGER*-*PROTEIN8* (*ZFP8*), meristem regulator *BAM2* and RuBisCo subunit *DEG24* were not differentially expressed in *RAP2*.*3*-transformed protoplasts. Specific activation and repression of the *TCL2* and *MYC3* promoter regions respectively was additionally confirmed through transactivation assays using RAP2.3 (**Fig 6b)**. Significant promoter activation in the presence of RAP2.3 was not detected for *ZFP8, DEG24,PSY1* and *BAM2*. While we were unable to detect *STM* transcripts in either control or *RAP2*.*3* transformed mesophyll protoplasts, likely due to strong epigenetic silencing of STM in developed leaves^31^, transactivation assays revealed that RAP2.3 could activate the *STM* promoter (**Fig.6c**). Promoter dissection study of *STM* revealed that a 450 bp region upstream of the translation start site is sufficient for its activation by both RAP2.3 and RAP2.12 (Supplemental data **Fig. S3**). Additionally, HRE2 and RAP2.2, but not HRE1, were able to activate this region (**Fig.6d**). In summary, our data shows that regulators of cell identity responsible for shaping leaf development are targeted by hypoxia transcription factors, linking oxygen distribution with developmental processes (**Fig.7**).

**Figure 6.**
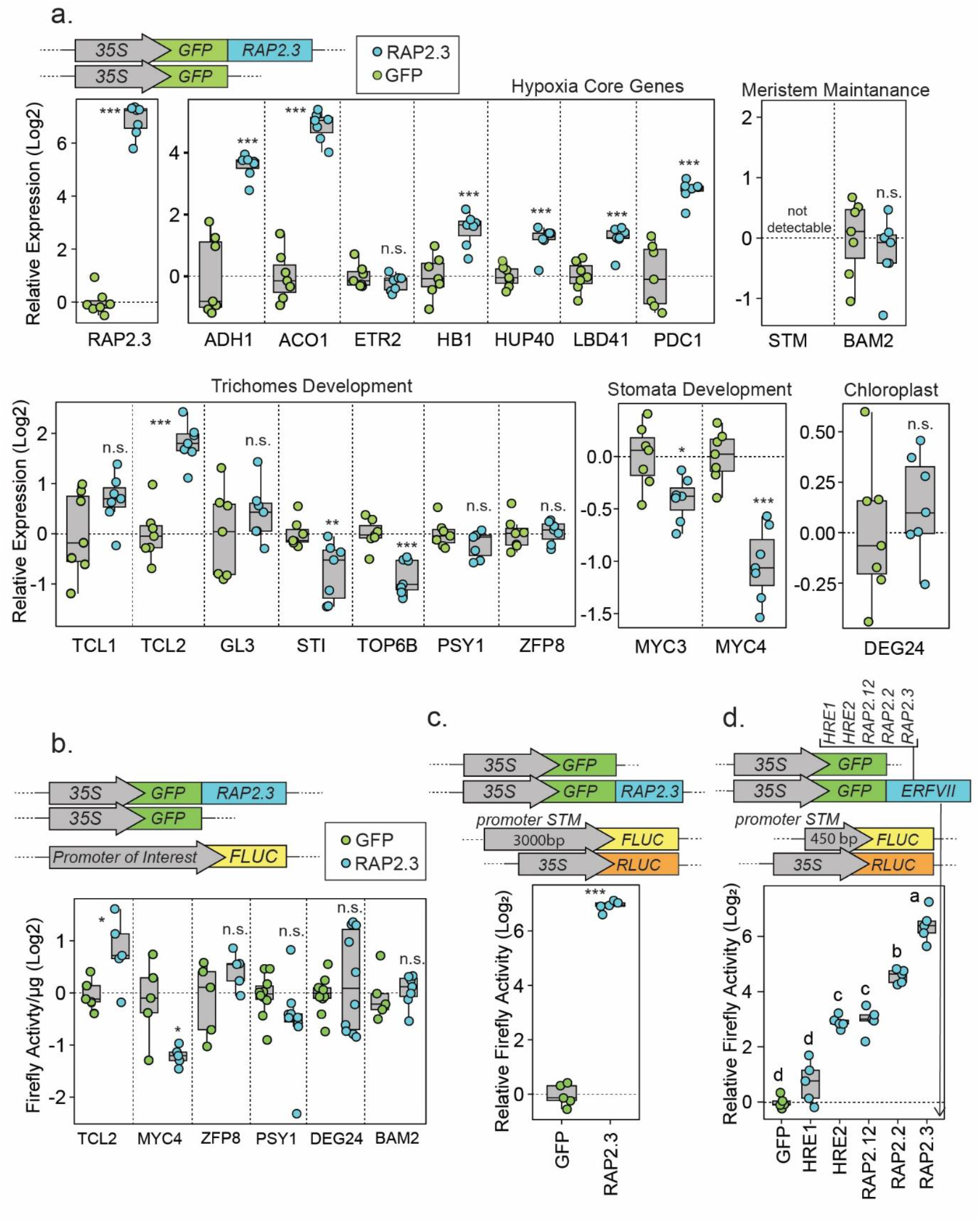
Molecular mechanisms underpinning cellular leaf traits are targets of oxygen sensing. **(a)** RT-qPCR on mesophyll protoplasts transformed with either *35S:GFP* (green) or *35S:GFP-RAP2*.*3* (cyan). Tested genes are grouped by the process they are associated with. Plots show relative expression on a log_2_ scale normalized to the relative expression of the *35S:GFP* protoplasts. **(b)** Transactivation assay testing the ability of GFP-RAP2.3 to activate or repress the promoter regions of candidate leaf development genes. Plots shows Firefly luciferase (FLUC) activity normalized on total ug of protein. **(c)** Transactivation assay on the STM promoter region to test the ability of GFP-RAP2.3 to induce *promSTM*. Plot shows relative Firefly Luciferase activity normalized to 35S:Renilla luciferase (RLUC). **(d)** Ability of the different *Arabidopsis thaliana* ERFVII to transactivate the 450 bp terminal region of the *STM* promoter. Plot shows relative Firefly Luciferase activity normalized to 35S: Renilla luciferase (RLUC). In **(a), (b), (c)** statistical differences are tested via Student or Welsh t-test, with asterisks indicating statically different groups (* p value < 0.05, ** p value < 0.01, *** p value < 0.001). In **(d)** statistical differences are assayed via one-way Anova and Tuckey post-hoc test for multiple comparisons, with letters indicating statistically different groups (p value < 0.05).

**Figure 7.**
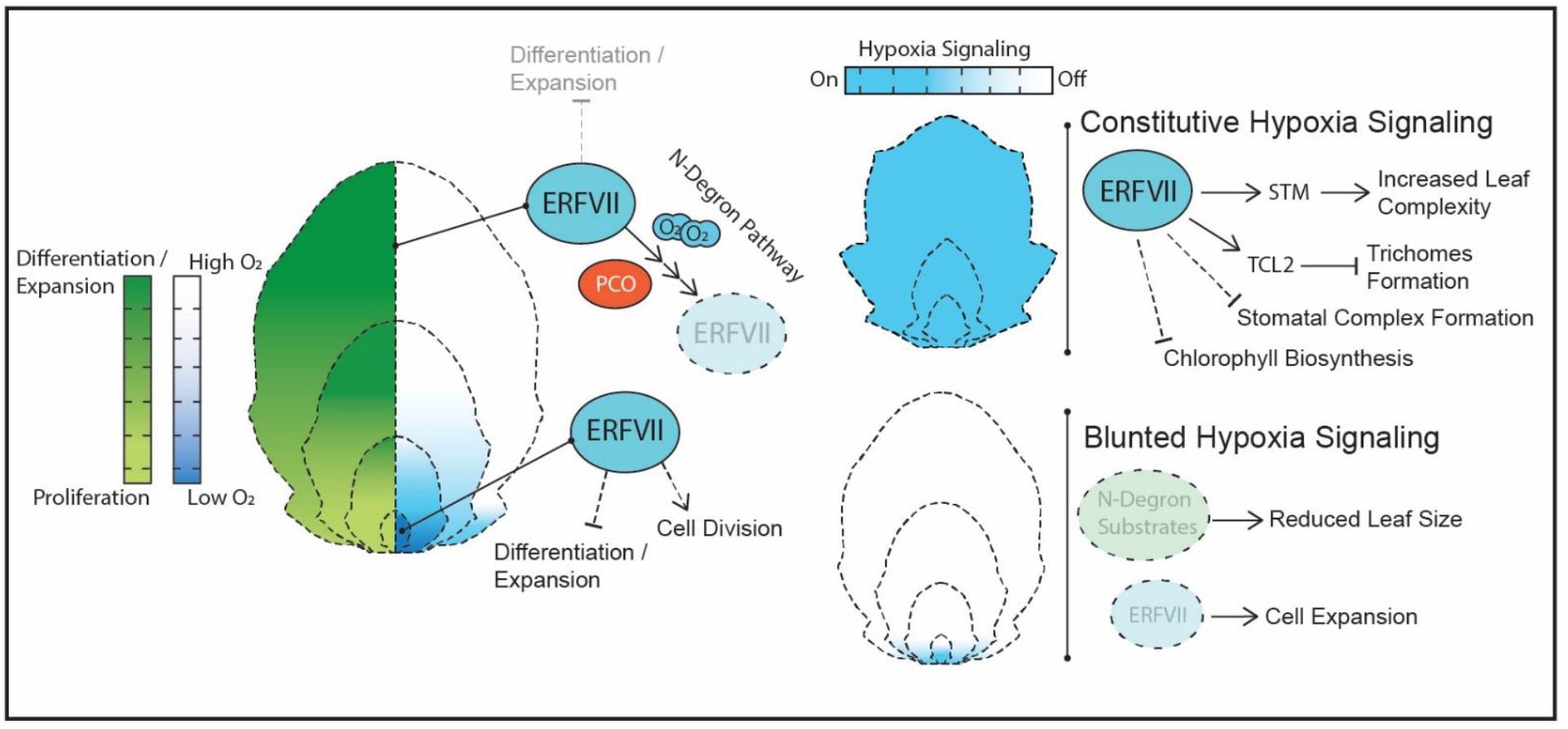
Proposed interplay of oxygen sensing and leaf development. Hypothetical model showing how progressive oxygenation during leaf development interplays with leaf morphology, cell expansion and fate acquisition. Initial and leaf proximal hypoxia promotes ERFVII stability, regulating cell expansion and cell division. Distal-to-proximal oxygenation destabilizes ERFVII via the PCO branch of the N-degron pathway mediating cellular differentiation and expansion. Constitutive hypoxia signaling or blunting of hypoxic responses interferes with leaf development as shown on the right.

## Discussion

Our study revealed that a spatiotemporal oxygenation pattern in leaves shapes specific aspects of leaf morphogenesis. We showed that on the whole-organ level, leaf primordia maintain a hypoxic status as they escape the meristem rather than abruptly oxygenating once they grow out of the shoot apex region. Interestingly, this hypoxic signature is maintained well beyond the very early stages of development, and is then progressively confined towards the proximal side of the leaf blade, creating an organ-wide gradient of oxygen-signaling. What features of the developing leaves are responsible for these oxygen gradients remain to be discovered. One factor may be spatial position: as leaves develop, they extend outward from the apex and re-establish contact with air, thereby facilitating gas exchange. We also observed an inverse relationship between chlorophyll fluorescence and the activation of hypoxia signaling in developing leaves. While this could reflect oxygenation of the tissue through photosynthesis, it may also instead represent negative regulation of chlorophyll biosynthesis by the oxygen-sensing machinery, as demonstrated here and previously reported in hypocotyls ^32^. Leaves were previously shown to experience cyclic fluctuations in oxygen content, due to cellular respiration at night and photosynthesis during the day ^33^. Although our measurements were conducted in the middle of the day, it remains to be determined whether similar fluctuations also occur during the early leaf stages examined here, prior to the establishment of a functional photosynthetic apparatus.

The initial hypoxic phase followed by distal-proximal gradual oxygenation does not appear to be merely a byproduct of leaf growth, but rather it appears to have been co-opted to regulate developmental processes. Indeed, prolonging the hypoxic phase through artificial depletion of available atmospheric oxygen led to an increased serration depth in fully mature leaves. Complementarily, both raising the oxygen level in the surrounding air to 40% or over expressing *PCO4* in developing leaves led to a significant decrease in final leaf size, suggesting that the initial hypoxic phase positively affects overall growth potential. These initial experiments led us to speculate a role for molecular oxygen in leaf development, although hyperoxic treatments may also affect energy conversion and lead to ROS accumulation, which can affect leaf development ^34^. Support for a signaling function of oxygen distribution was provided by the strong leaf phenotype of the *4pco* mutant, which leads to stabilization of ERFVII and induction of *HRG*s. The rescue of increased complexity of *4pco* by *4pco3rap* strongly points towards ectopic stabilization of ERFVII as the main contributor to the severe *4pco* phenotype. Interestingly, the *erfvii* mutant did not display leaf morphological changes when compared to wildtypes, but did exhibit a marked increase in epidermal cell size, suggesting an additional role for ERFVII in inhibiting cellular expansion. Alternatively, the increased cell size might be a secondary compensatory mechanism triggered by an impairment in cell division in the mutant. Destabilization of PCO substrates through hyperoxia treatments or *PCO4* overexpression did result in smaller leaves, suggesting that other Cys-initiating proteins may be involved in regulating cell expansion and final leaf size. Notably, one such target, VRN2 has previously been shown to regulate cell expansion and rosette size ^8^.

Our *4pco* transcriptome dataset showed misregulation of key factors which could be connected to specific leaf development processes, suggesting a broad impact of oxygen sensing on leaf development. Furthermore, these transcriptional deviations were associated with actual phenotypes in *4pco*. First, stomatal production is impaired in *4pco*, consistent with a down regulation of *SOL1* and an upregulation of negative regulators of stomata development *MYC4* and *MPK3*. Surprisingly, we found that *MYC3* and *MYC4* were actually downregulated by RAP2.3 overexpression, indicating that their induction in *4pco* might be due to other N-degron pathway substrates. Second, trichome density is severely decreased in *4pco* and partially rescued in the *4pco-3rap*, suggesting additional regulation independent of RAP2.2/2.3/2.12. The upregulation of *TCL2*, a competitor of *GLABRA1* during the formation of the trichome initiation complex, is consistent with the reduction of adaxial trichomes in *4pco*. Moreover, *STI* and *TOP6B*, both necessary for trichome patterning and branching, are downregulated in both *4pco* leaves and RAP2.3-overexpressing protoplasts. Third, while lobeyness of epidermal cells increases towards the distal side of the leaf in wildtype plants, it remains stably low in *4pco* leaves. We found upregulation of *STM* in 4pco leaves, and transactivation of its promoter by RAP2.3. Ectopic expression of *STM* was previously shown to increase leaf complexity and delay epidermal lobeyness and thus differentiation^19^. Fourth, many chlorophyll biosynthesis and chloroplast organization genes are misregulated in *4pco*, and chlorophyll biosynthesis is severely impaired in *4pco*. Taken together, this data suggests an overall impairment in cell fate specification and cellular differentiation when hypoxia signaling is not switched off, indicating that the signals originated during normal distal-to-proximal oxygenation of the leaf are required for acquisition of specialized cell-types. Our results further indicate that multiple developmental regulators act downstream of the oxygen-sensing machinery, potentially connecting oxygen distribution directly to leaf morphogenesis. Additional studies will be required to fully elucidate the underlying mechanisms.

Taken together, our study shows that leaves initiate in hypoxia, and progressively become oxygenated from tip to base as they leave the shoot apex. Permanence of hypoxia during leaf development promotes increases leaf complexity, and inhibits the acquisition of specialized cell-types in an ERFVII depending manner, while the initial hypoxic state instead appears to be involved in regulation of cell expansion. A more complex leaf shape has been shown to increase gas exchange in plant species that display heterophylly ^35^. Interestingly, in *Hygrophila difformis* and *Rorippa aquatica* heterophylly is mediated through a delay in differentiation via *STM*, which we also found induced in *4pco* ^36,37^. It is tempting to speculate that the increased leaf complexity induced by persistent hypoxia during leaf development in Arabidopsis is mechanistically reminiscent of more sophisticated heterophyllic adaptations of aquatic and amphibious plants to their wet environments. Hence, while here we found that oxygen distribution acts as an endogenous cue to regulate leaf morphology, this also lays the groundwork to investigate how environmental oxygen availability acts as a signaling factor influencing leaf shape.

## Materials & Methods

### Plant Materials

*Arabidopsis thaliana* Columbia-0 seeds were used as a wild-type ecotype. The *pPCO1:GUS-GFP* line and the *4pco* and *erfvii* mutants were previously described^2,38^. The *4pco/3rap* lines used in this study was newly generated through CRISPR-Cas9 gene editing *4pco* background. The *RPS5a:XVE/pLexA:PCO4-mCherry* was generated as described in the cloning section.

### Plants Growth Conditions

All plants used in this study were grown on soil in short day conditions (8h light, 16h darkness, 21°C, 70% RH, 200 µmol/m^2^/s light intensity). Seeds were vernalized for 7 days at 4°C degrees prior to growth.

### Vibratome Sectioning of Shoot Apical Meristems

Mature leaves from *Arabidopsis thaliana* 4-weeks old plants were removed in order to isolate the shoot apex. Samples were then embedded in 4% agarose and sectioned longitudinally using a Leica VT1000S vibratome. Average sections were 120 µm thick.

### Microsensor profiling

Oxygen microprofiling was conducted on emerging leaves of 4-week-old Arabidopsis plants using a Clark-type microsensor with a 10 μm tip (Unisense). The sensor was linked to a pA meter (Unisense) and mounted on a motorized micromanipulator (MM33, Unisense). Measurements were taken in 10 μm steps, beginning outside the leaf surface and advancing until the desired tissue region was reached. Before use, the microsensor was calibrated under two reference conditions: 0% O_2_, achieved with a sodium ascorbate solution (50 mL sodium ascorbate in 50 mL 0.1 M NaOH), and 21% O_2_, obtained by equilibrating medium with air. A dissection microscope on a boom stand (Zeiss Stemi 305) was used to precisely position the probe above the tissue. All profiling experiments were carried out at 20 °C under low light.

### Histological Staining

Developing leaves of *pPCO1:GUS-GFP* were separated from the meristem with a sharp razorblade and used for histochemical GUS staining. Leaves were fixed for 1h in 90% Acetone on ice, washed twice with water and immersed in GUS staining solution (100 mM buffer phosphate, 0.1% Triton X-100, EDTA pH 8 10 mM, potassium ferrocyanide 0.5 mM, potassium ferricyanide 0.5 mM, X-Gluc 200 mM) for 4 hours. Leaves were then cleared in several washes of 70% (v/v) ethanol and imaged using a Leica M205 FCA stereomicroscope equipped with a Leica dfc7000t camera.

### GFP and Chlorophyll Imaging and Quantification

Excised developing leaves were imaged using a Zeiss Apotome 3 fluorescence microscope with a band-pass filter for GFP (excitation: 450-490, emission: 500-550) and Chlorophyll (excitation: 625-655, emission: 665-715). Signal intensity across the leaf blade was quantified in Fiji. To improve visualization, the Royal look-up-table was applied on a set of representative images in figures 1c and 5e.

### Controlled gas treatments

Plants were exposed to low-oxygen or high oxygen conditions inside plexiglass chambers that were continuously supplied with a humidified gas mixture of oxygen and nitrogen, at the ratios specified in the Results section. 440 ppm CO_2_ was added to prevent carbon limitation. During hypoxia treatments, the chambers were kept in under normal light conditions to avoid dark responses. Control plants were kept in the light for the same duration and flushed with air (21% O_2_) v/v.

### Construct Design and Induction in plants

The *pRPS5A:XVE-pLEXA:PCO4-mCherry* was generated using Green Gate cloning technology ^39^. The *PCO4* coding sequence was cloned from Col-0 complementary DNA (cDNA) without its endogenous STOP codon and placed upstream of a *mCherry* fluorescent protein and downstream of a *LexA* promoter in a *pGGN000* intermediate plasmid. A *pRPS5-XVE* module was assembled in a *pGGM000* intermediate module. The two intermediate modules were then combined in a *pGGZ001* to form a single construct with two transcriptional units and a Hygromycin plant resistance. The construct was induced by spraying soil-grown plants with a solution of 50 µM β - estradiol (dissolved in DMSO). Plant were sprayed every 48h over the induction period and activation of *PCO4* was monitored via imaging of mCherry fluorescence.

### Generation of CRISPR/Cas9 lines

CRISPR/Cas9 guide constructs were generated by assembling synthetic DNA fragments into a modular expression cassette. The cassette included a U6 promoter for RNA polymerase III–driven transcription, a Glycine-tRNA sequence to facilitate the multiplexing and processing of multiple guide RNAs, the guide RNA sequences, and the conserved sgRNA scaffold required for Cas9 recognition. Transcriptional termination was ensured by a poly-T stretch42. The entire cassette was flanked by *attB* recombination sites to enable Gateway® cloning into binary destination vectors for plant transformation. Guide sequences (gRNAs) used in this study can be found on table S1.

### Leaf Phenotyping

Fully developed leaves were cut at the site of the petiole, flattened with tape against a sheet of paper and scanned using an Epson V850 Pro scanner. Binary leaf silhouettes were generated in Fiji and imported into R using the *magick* and *imager* packages. Leaf contours were extracted from the images and resampled to a fixed number of equidistant points along the perimeter. Leaf area and perimeter were calculated directly from the extracted contours using the shoelace formula. Elliptical Fourier Analysis (EFA) was then applied to the resampled contours to reconstruct smooth leaf outlines for subsequent shape descriptors. The convex hull of each leaf was determined using the base R function chull(), and leaf solidity was calculated as the ratio of leaf area to convex hull area. Average serration depth, representing the mean shortest distance from contour points to the convex hull, was used to quantify lobing. The first two proximal serrations were used for this calculation. Principal Component Analysis (PCA) of the reconstructed coordinates was used to define the major and minor axes.

### Cellular Phenotyping

Fully developed leaves were washed and cleared by an overnight incubation in 70% v/v/ Ethanol, followed by an overnight incubation in Hoyer’s solution (Chloral hydrate/water/glycerol 8 W: 2 V: 1 V)^40^.Distal and proximal regions along the main leaf axis were dissected and the abaxial epidermis was imaged using Zeiss Apotome 3 equipped with a DIC module. Cell contours for segmentation and convex hulls were produced in Fiji. Area and perimeter of both segmented cells and their convex hulls were computed in Fiji. Lobeyness was calculated as the ratio between the perimeter of each cell and the perimeter of their convex hull. In order to remove stomata and stomatal lineage cells from the calculations, only cells with an area larger than 600 um were considered. For visualization, epidermal cells were coloured based on either surface area or lobeyness. For lobeyness, colour gradients were assigned using the ROI Color Coder macro^41^. Counting of stomatal complexes was also performed on the same DIC images.

### Trichome Quantification

Brightfield images of leaves 7 from different oxygen sensing mutants were used for quantifying adaxial trichomes in Fiji. Images were acquired using a Leica M205 FCA stereomicroscope equipped with a Leica dfc7000t camera.

### RNA-Seq preparation and analysis

Leaf 7 of the relevant developmental stages were harvested from developing Col-0 and *4pco* plants. Approximately 10 leaves were pooled for each biological replicate. Total RNA was extracted with a Quick-RNA Miniprep kit from Zymo Research, according to manufacturer instructions. Genomic DNA removal was performed with on-column DNAse I treatment. Both mRNA library preparation (poly-A enrichment) and sequencing were performed by Novogene. Raw sequencing quality was assessed via FastQC. Trimmed reads produced via TrimGalore were mapped to the TAIR10 Genome using Hisat2. Absolute counts were obtained through featureCounts. For each gene, raw counts were modeled using a generalized linear model with a negative binomial distribution. The design formula (∼ Genotype + Stage + Genotype:Stage) allowed testing for baseline differences between genotypes, temporal changes within the reference genotype, and whether temporal responses differed between genotypes (interaction). Statistical significance of each coefficient was assessed using Wald tests adjusted by Benjamini-Hochberg procedure for multiple testing. An adjusted p value of 0.01 was used as a cutoff to identify genes with a significant genotype, developmental stage or interaction effect. In order to identify candidate genes to connect low oxygen sensing and leaf development, we cross referenced the list of genes with a significant genotype or interaction effect with a list of genes associated with leaf development GO categories (full list on table S3).

### Protoplast Isolation and Transformation

Mesophyll protoplasts were isolated from leaves of 4-week-old *Arabidopsis thaliana* Col-0 plants using the Tape-Arabidopsis Sandwich method. The abaxial epidermis was removed, and the exposed mesophyll tissue was digested for 2 h in the dark in an enzyme solution containing 1.5% cellulase R10, 0.4% macerozyme R10, 0.4 M mannitol, 20 mM KCl, and 20 mM MES (pH 5.7). Prior to use, the enzyme solution was incubated at 55 °C for 10 min, cooled to room temperature, and supplemented with 0.1% BSA and 10 mM CaCl_2_.Released protoplasts were collected in 40-mL tubes and pelleted at 100 × g for 2 min at 4 °C. The supernatant was removed, and protoplasts were washed once with an equal volume of W5 solution (154 mM NaCl, 125 mM CaCl_2_, 5 mM KCl, 2 mM MES, pH 5.7). Cells were kept on ice, in the dark, for 20 min and then pelleted again at 100 × g for 1 min at 4 °C. The supernatant was removed, and the pellet was resuspended in MMG solution (0.4 M mannitol, 15 mM MgCl_2_, 4 mM MES, pH 5.7). For transformation, 100 µL of protoplast suspension was mixed with the desired plasmids and 100 µL of PEG solution (prepared by dissolving 4 g PEG4000 in 3 mL H_2_O, 2.5 mL 0.8 M mannitol, and 1 mL CaCl_2_; final volume 10 mL). After 5 min incubation in the dark, transformation was stopped by adding 440 µL of W5 solution. Protoplasts were pelleted at 100 × g for 2 min, the supernatant was removed, and cells were resuspended in 1 mL of WI solution (0.5 M mannitol, 20 mM KCl, 4 mM MES, pH 5.7, 10 mM glucose). Protoplasts were incubated overnight in the dark prior to use in transactivation assays or RNA extraction.

### Protoplast RNA extraction and qPCR

Total RNA was extracted from transformed mesophyll protoplasts. Five independent transformation aliquots were pooled for each assayed biological replicate. RNA was extracted with Quick-RNA Miniprep kit (Zymo Research), according to manufacturer instructions. Genomic DNA was removed via on-column DNAse I digestion. Following extraction, complementary cDNA was synthesized via RevertAid First Strand cDNA Synthesis Kit (ThermoFisher Scientific). RT-qPCR was set up with SYBR Green Real-Time Master Mix and run 10 ng of cDNA. Primers used in this study can be found on table S1.

### Transactivation assays

Transactivation assays were carried out using a dual-luciferase system using *Renilla reniformis* and *Photinus pyralis* luciferases. Target promoters were cloned from genomic DNA, and then cloned into *pDONR207* (Thermo Fisher Scientific) using a BP reaction, and subsequently recombined into the *PGWL7* vector (Hellens et al., 2005) using LR Clonase II (Thermo Fisher Scientific) to create the reporter construct promoter:PpLuc. A 35S:Renilla, or total protein content was used for normalization. To test the impact of RAP2.3 on each promoter, effector plasmids were prepared by transferring the coding sequences of *RAP2*.*3* from *pENTR-D/TOPO* into *p2GW7* (Karimi et al., 2002). Mesophyll protoplasts were isolated as described above and transfected with 5 μg of each plasmid. Following a 16-hour incubation in WI medium, proteins were extracted using 30 μL of 1× Passive Lysis Buffer (Promega). Luciferase activities were quantified using the Dual-Luciferase Reporter Assay kit (Promega) and a GloMax 20/20 luminometer (Promega). Total protein content was measured using a colorimetric Bradford assay.

## Supplemental Information

**Supplementary Figure S1**

CRISPR/Cas9-induced mutations in RAP2.2, RAP2.3 and RAP2.12 genes

**Supplementary Figure S2**

Chlorophyll development over leaf growth in *Arabidopsis thaliana*

**Supplementary Figure S3**

Dissection of the STM promoter region as a target of RAP2.12 activation

**Supplementary Table 1**

Primers and guideRNAs used in this study.

**Supplementary Table 2**

Description of RPS5:PCO4 Inducible Construct.

**Supplementary Table 3**

GO Terms used for Transcriptome Investigation.

## Acknowledgements

The authors acknowledge the funding provided for DAW, GP, and VV by the European Research Council (ERC) grant 101077812, and DAW and VV by the Netherlands Organization for Scientific Research (NWO), VIDI grant VI.Vidi.213.055. LJ was funded by NWO VIDI grant number VI.Vidi.193.104 granted to Kaisa Kajala (Utrecht University). FL, LDC and VS were funded by the ERC grant 101001320 and FL and VS by UKRI Biotechnology and Biological Science Research Council grant application APP23352

## Author contributions

Experiments were conceived by DW, GP and FL and performed by GP, VS, VV, KK, SB and LDC. Data analysis was performed by GP, DW and LJ. The manuscript was written by GP and DW and commented on by FL and LJ. All authors approved the final version of the manuscript.

---

**Figure S1.**
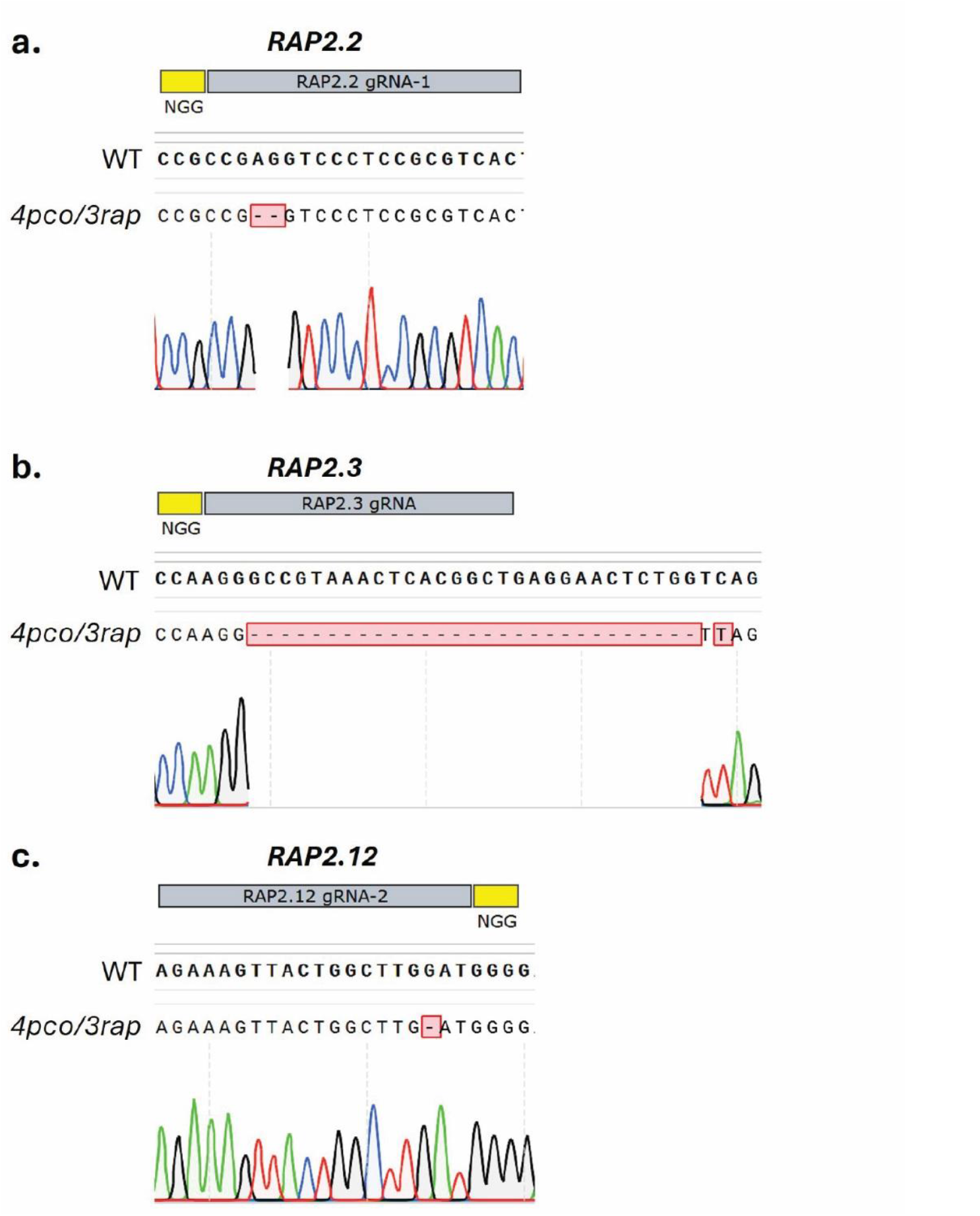
CRISPR/Cas9-induced mutations in RAP2.2, RAP2.3 and RAP2.12 genes. Sequence chromatograms showing representative mutations generated at target sites of RAP2.2, RAP2.3, and RAP2.12 using the indicated gRNAs. **(a)** RAP2.2 gRNA-1: 1-bp deletion within the target sequence. **(b)** RAP2.3 gRNA: large deletion spanning the protospacer region. **(c)** RAP2.12 gRNA-2: 1-bp insertion at the target site. Target sequences are shown above chromatograms, with NGG PAM motifs highlighted. Red boxes indicate the position and type of mutation.

**Figure S2.**
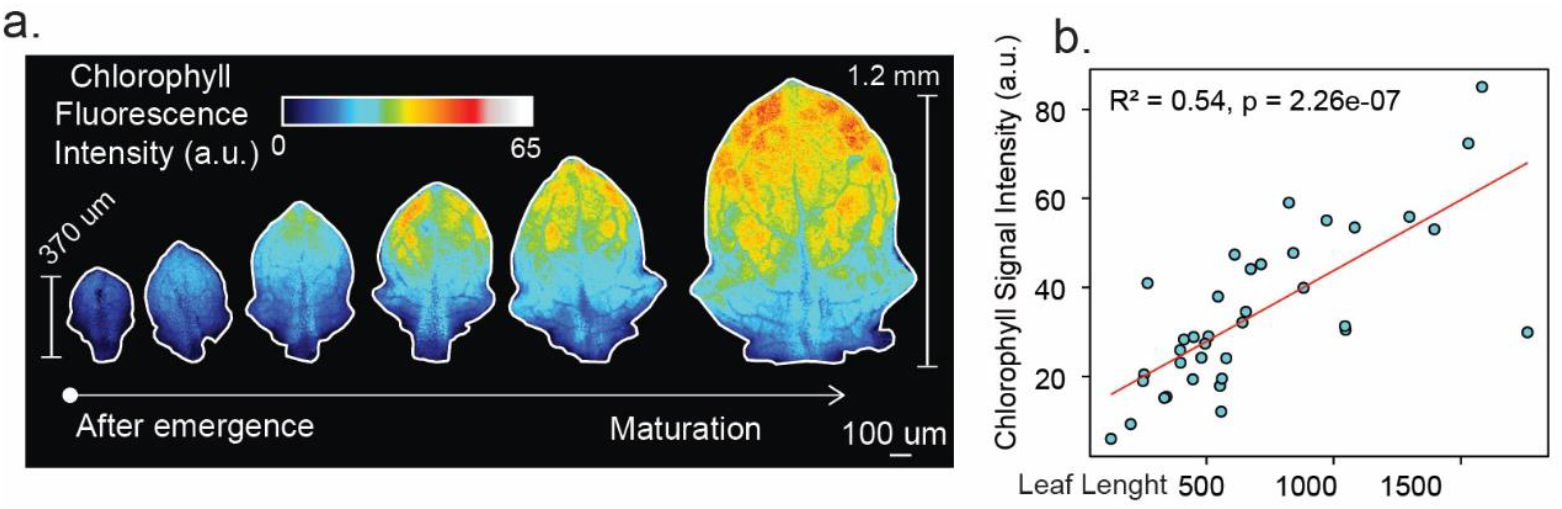
Chlorophyll development over leaf growth in *Arabidopsis thaliana*. **(a)** Imaging of Chlorophyll autofluorescence across *promPCO1:GFP* developing leaves 7 ± 1 shown in figure 1c. Signal visualized with royal look-up-table. **(b)** Quantification of Chlorophyll autofluorescence in *promPCO1:GFP* developing leaves in relation to the size of their major axis. Red line represents the linear regression model fitted to the Chlorophyll signal as a function of length. Model fit statistics (R^2^ and p-value for the slope) are indicated on the plot.

**Figure S3.**
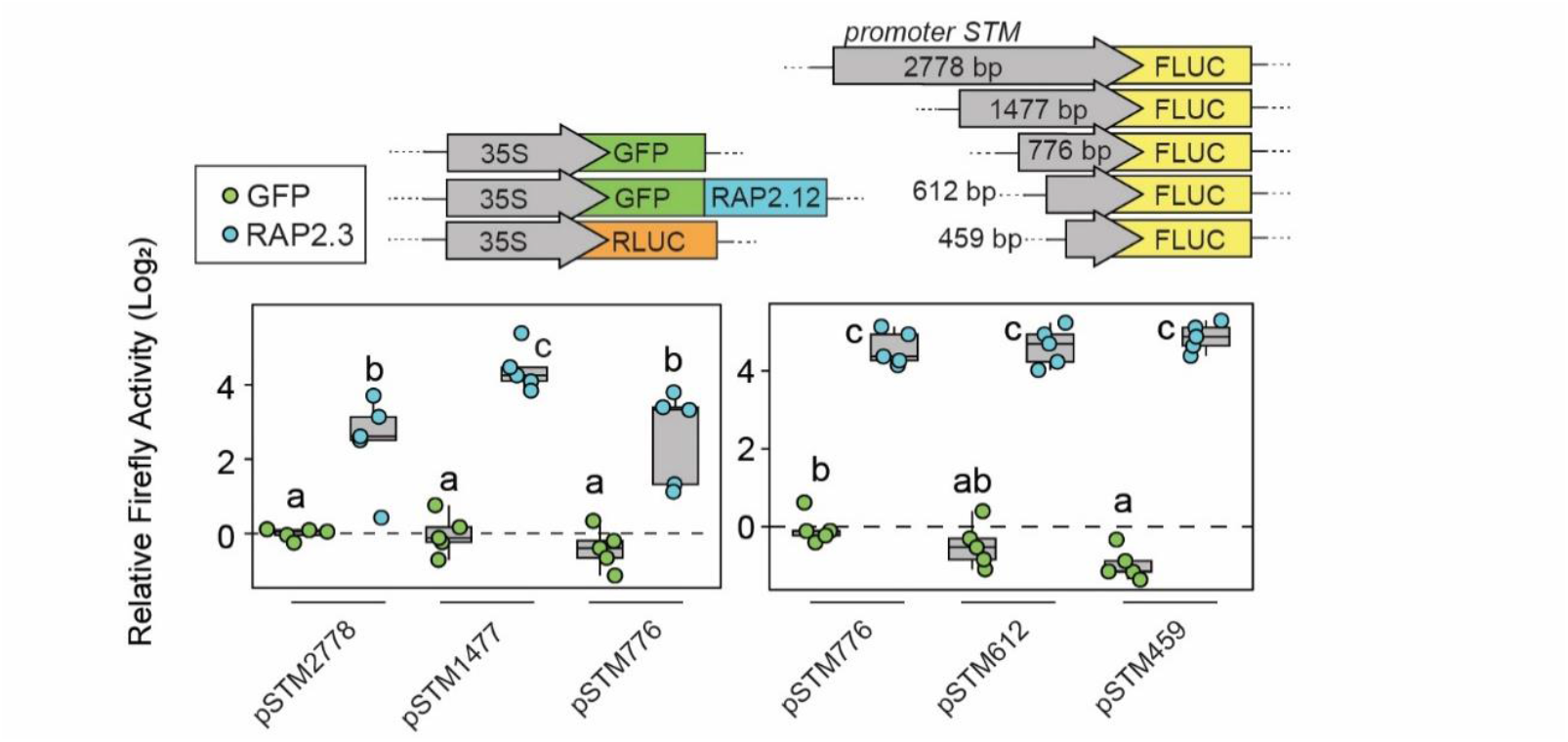
Dissection of the STM promoter region as a target of RAP2.12 activation. Transactivation assay of GFP-RAP2.12 on progressively shorter regions of the *STM* promoter region. Plot shows relative Firefly Luciferase activity normalized to 35S:Renilla luciferase (RLUC). Statistical differences are assayed via two-way Anova and Tuckey post-hoc test for multiple comparisons, with letters indicating statistically different groups (p value < 0.05).

