## Supplementary Table 1 for "Progressive oxygenation of developing leaves directs morphogenesis"

**Supplementary Table S1. List of primers used in this study**

**Cloning of promoter Regions:**

| **Primer name** | **5’-3’ sequence** | **AGI-code** |
| --- | --- | --- |
| pTCL2_Fw_gw | GGGGACAAGTTTGTACAAAAAAGCAGGCTGATGCATTACCAAAATCACTCCACC | at2g30424 |
| pTCL2_Rv_gw | GGGGACCACTTTGTACAAGAAAGCTGGGTCCAGTGGTATTTGAAAGTGAAAGAGG | at2g30424 |
| pMYC4_Fw_gw | GGGGACAAGTTTGTACAAAAAAGCAGGCTGCTTATCATTTTCACCTTTTCTTAAACC | at4g17880 |
| pMYC4_Rv_gw | GGGGACCACTTTGTACAAGAAAGCTGGGTGGAGACGTAACAGTTCTCTGACGTAG | at4g17880 |
| pDEG24Fw | GGGGACAAGTTTGTACAAAAAAGCAGGCTTCGCAACACTAAGCCTACGA | AT5G38420 |
| pDEG24Rv | GGGGACCACTTTGTACAAGAAAGCTGGGTGGAGAACATAGAGGAAGCCAGTAC | AT5G38420 |
| pBAM2Fw | GGGGACAAGTTTGTACAAAAAAGCAGGCTTGTTTGTGTCTCTCCCAATTGA | AT3G49670 |
| pBAM2Rv | GGGGACCACTTTGTACAAGAAAGCTGAGGTTAGCGACGGTGAAGGAATGA | AT3G49670 |
| pPSY1Fw | GGGGACAAGTTTGTACAAAAAAGCAGGCTGCCGGTGGATATGACTCACT | AT5G58650 |
| pPSY1Rv | GGGGACCACTTTGTACAAGAAAGCTGGGTTTGCGGGCTAGAGAAGATGT | AT5G58650 |
| pZFP8Fw | GGGGACAAGTTTGTACAAAAAAGCAGGCTCGTGGACTCGTCATGTGAAG | AT2G41940 |
| pZFP8Rv | GGGGACCACTTTGTACAAGAAAGCTGGGTCTCTCGTCCGTTGGTTTCG | AT2G41940 |
| pSTM2778bp_Fw | CACCCTGGACTTTCCGAGGCACATT | AT1G62360 |
| pSTM459bp | CACCAGAACCCACAGATCACACA | AT1G62360 |
| pSTM_Rv | CTTCTCTTTCTCTCACTAGTA | AT1G62360 |

**Primers for RT-qPCR**

| **Primer name** | **5’-3’ sequence** | **AGI-code** |
| --- | --- | --- |
| qTCL2_FW | ACCGCAGAGGTAAATAGTGTGAAA | AT2G30424 |
| qTCL2_RV | TAAATCCCACCTATCACCAACAAG | AT2G30424 |
| qSTI_FW | CGACTTTGCTTCAGCTTGGG | AT2G02480 |
| qSTI_RV | TGGGTCATCGTCAGTTGCTC | AT2G02480 |
| qZFP8_FW | CCACCGACAATCAACGGAAG | AT2G41940 |
| qZFP8_RV | TCATGCGACCTAAACGCTGA | AT2G41940 |
| qPSY1_FW | CTGTTTCCGTTTCAGGTGGG | AT5G58650 |
| qPSY1_RV | GGGTCACCGTAGTCCTCAAC | AT5G58650 |
| qTOP6B_FW | GGTAAGGGGATGCCACATGA | AT3G20780 |
| qTOP6B_RV | CCATCTTTGCACCCAGACCA | AT3G20780 |
| qDEG24_FW | TTCATTGCCTACAAGCCCCC | AT5G38420 |
| qDEG24_RV | ATGGGTTCCAGAATAATCAACGC | AT5G38420 |
| qGL3_FW | GTGTGGTTTCATGGGATATAGGG | AT5G41315 |
| qGL3_RV | CGAACTGAAACTGCGAGGTGT | AT5G41315 |
| qTCL1_FW | AGAAGAGTGGTGGGACGTGA | AT2G30432 |
| qTCL1_RV | ACTTTTGCGTCATTTGTGGGA | AT2G30432 |
| qMYC3_FW | TCATCCCGGTGCTAGGTTCA | AT5G46760 |
| qMYC3_RV | GCTCCCCATCTTCACCGTAG | AT5G46760 |
| qMYC4_FW | GCCGTGTTCTGGCAATCATC | AT4G17880 |
| qMYC4_RV | ACCATCTCCCCAACCTAACAA | AT4G17880 |
| qBAM2_FW2 | ACGTGACTGAGAAAGCTCCG | AT3G49670 |
| qBAM2_RV2 | CAAACCCAATCGCCGGAAAG | AT3G49670 |
| qSTM_FW | ACCTTCCTCTTTCTCCGGTTATGG | AT1G62360 |
| qSTM_RV | GCGCAAGAGCTGTCCTTTAAGC | AT1G62360 |
| qUBQ10_F | GGCCTTGTATAATCCCTGATGAATAAG | AT4G05320 |
| qUBQ10_R | AAAGAGATAACAGGAACGGAAACATAGT | AT4G05320 |
| qETR2FW | TGTTAGATTCTCCGGCGGCTATG | AT3G23150 |
| qETR2REV | TTCCCATGAATCAACTGCACCAC | AT3G23150 |
| qHUP40 _FWD | GAAACTTGAGTGCGAGTGTG | AT4G24110 |
| qHUP40 _REV | CTCAAACCCAATCTTTTGCT | AT4G24110 |
| qPDC1FW | GACGCCATTCATAACGGTGAAG | AT4G33070 |
| qPDC1RV | GGATTGGGAGGACGGCTGT | AT4G33070 |
| qADH1FW | TCACTGTTGATAATGTCTACCACCG | AT1G77120 |
| qADH1RV | CATGGCCGAAGATACGTGGA | AT1G77120 |
| qLBD41_FW | TGAAGCGCAAGCTAACGCA | AT3G02550 |
| qLBD41_RV | ATCCCAGGACGAAGGTGATTG | AT3G02550 |
| qHB1_FW | TTTGAGGTGGCCAAGTATGCA | AT2G16060 |
| qHB1_RV | TGATCATAAGCCTGACCCCAA | AT2G16060 |
| qACO1FW | ACCAGTCAGAGATGGTCAAGGC | AT4G35830 |
| qACO1REV | TCATCCATCGTCTTGCTGAGTTCC | AT4G35830 |
| qRAP2.3FW | AACTCACGGCTGAGGAACTCTG | AT3G16770 |
| qRAP2.3REV | ACGTTAACTTGGTTGGTGGGATGG | AT3G16770 |

**Guides for RAP2.2/2.3/2.12 CRISPR-Cas9**

| RAP2.12_1 | *TGCGATGATGATTTCGACGT* | AT1G53910 |
| --- | --- | --- |
| RAP2.12_2 | *AGAAAGTTACTGGCTTGGAT* | AT1G53910 |
| RAP2.2_1 | *GTGACGCGGAGGGACCTCGG* | AT3G14230 |
| RAP2.2_2 | *ATTTCGAAGCTGATTTCCAA* | AT3G14230 |
| RAP2.3 | *AGCCGTGAGTTTACGGCCCT* | AT3G16770 |
