## Supplementary Table 2 for "Progressive oxygenation of developing leaves directs morphogenesis"

**Supplementary Table S2. Sequence of modules used to achieve PCO4 inducible overexpression:**

Full plasmid name:

*promRPS5: Ω-XVE-tUB10;pLEXA:Ω-PCO4-mCherry-tUB10-HygR* in *pGGZ003* backbone.

Sequences for *promRPS5a* (pGGA012), *Ω viral 5’UTR* (pGGB002), *UBQ10 terminator* (*tUB10*) (pGGE009), *mCherry*-*linker* (pGGD003) and Hygromycin resistance (*HygR*) (pGGF005) are those described by Lampropoulos *et al*. in the Green Gate technology plasmid set ^1^. Sequences of custom modules follow.

XVE: atgaaagcgttaacggccaggcaacaagaggtgtttgatctcatccgtgatcacatcagccagacaggtatgccgccgacgcgtgcggaaatcgcgcagcgtttggggttccgttccccaaacgcggctgaagaacatctgaaggcgctggcacgcaaaggcgttattgaaattgtttccggcgcatcacgcgggattcgtctgttgcaggaagaggaagaagggttgccgctggtaggtcgtgtggctgccggtgaaccgtcgagcgcccccccgaccgatgtcagcctgggggacgagctccacttagacggcgaggacgtggcgatggcgcatgccgacgcgctagacgatttcgatctggacatgttgggggacggggattccccgggtccgggatttaccccccacgactccgccccctacggcgctctggatatggccgacttcgagtttgagcagatgtttaccgatgcccttggaattgacgagtacggtggggatccgtctgctggagacatgagagctgccaacctttggccaagcccgctcatgatcaaacgctctaagaagaacagcctggccttgtccctgacggccgaccagatggtcagtgccttgttggatgctgagccccccatactctattccgagtatgatcctaccagacccttcagtgaagcttcgatgatgggcttactgaccaacctggcagacagggagctggttcacatgatcaactgggcgaagagggtgccaggctttgtggatttgaccctccatgatcaggtccaccttctagaatgtgcctggctagagatcctgatgattggactcgtctggcgctccatggagcacccagggaagctactgtttgctcctaacttgctcttggacaggaaccagggaaaatgtgtagagggcatggtggagatcttcgacatgctgctggctacatcatctcggttccgcatgatgaatctgcagggagaggagtttgtgtgcctcaaatctattattttgcttaattctggagtgtacacatttctgtccagcaccctgaagtctctggaagagaaggaccatatccaccgagtcctggacaagatcacagacactttgatccacctgatggccaaggcaggcctgaccctgcagcagcagcaccagcggctggcccagctcctcctcatcctctcccacatcaggcacatgagtaacaaaggcatggagcatctgtacagcatgaagtgcaagaacgtggtgcccctctatgacctgctgctggagatgctggacgcccaccgcctacatgcgcccactagccgtggaggggcatccgtggaggagacggaccaaagccacttggccactgcgggctctacttcatcgcattccttgcaaaagtattacatcacgggggaggcagagggtttccctgccacagtctga

pLEXA:
agcttgggctgcaggtcgaggctaaaaaactaatcgcattatcatcccctcgacgtactgtacatataaccactggttttatatacagcagtactgtacatataaccactggttttatatacagcagtcgacgtactgtacatataaccactggttttatatacagcagtactgtacatataaccactggttttatatacagcagtcgaggtaagattagatatggatatgtatatggatatgtatatggtggtaatgccatgtaatatgctcgactctaggatcttcgcaagacccttcctctatataaggaagttcatttcatttggagaggacacgctgaagctagtc

PCO4: atgccttactttgctcagaggctttacaatacttgcaaggcgtctttctcttcagacggaccaataacagaagatgctctggagaaggttcgcaatgtcttggagaaaatcaagccatctgatgttggaattgagcaggatgctcaattggcacggtccaggtctggtcctctcaatgaacgcaatggaagtaatcagtctcccccagcaataaagtatcttcatttgcatgagtgtgacagtttctctataggaatcttctgtatgccaccttcttctatgatacctcttcataaccatccgggcatgaccgtgctaagcaagctcgtttatggttcaatgcatgtgaagtcatatgattggctagagcctcaactgaccgaaccagaggatccatcacaagcaagacctgctaaactggtgaaggatactgagatgacggctcaaagcccagtaacaacattatatccgaaaagtggtggcaacattcactgtttcaaagccatcacccattgtgctattcttgacatcttagctccaccttactcttcagagcatgatcggcattgcacttacttccggaaatccagaagagaagacttaccaggtgaattagaagtggatggagaagtggtcacagacgtgacatggcttgaggaatttcaaccgcctgatgactttgtgatacgacgaattccgtacagaggtcctgtcattagaacttga
